# Persistent DNA-break potential near telomeres increases initiation of meiotic recombination on short chromosomes

**DOI:** 10.1101/201889

**Authors:** Vijayalakshmi V. Subramanian, Xuan Zhu, Tovah E. Markowitz, Luis A. Vale-Silva, Pedro A. San-Segundo, Nancy M. Hollingsworth, Scott Keeney, Andreas Hochwagen

## Abstract

Faithful meiotic chromosome inheritance and fertility relies on the stimulation of meiotic crossover recombination by potentially genotoxic DNA double-strand breaks (DSBs). To avoid excessive damage, feedback mechanisms down-regulate DSBs on chromosomes that have successfully initiated crossover repair. In *Saccharomyces cerevisiae*, this regulation requires the removal of the conserved DSB-promoting protein Hop1/HORMAD during chromosome synapsis. Here, we identify privileged end-adjacent regions (EARs) spanning roughly 100 Kb near all telomeres that escape DSB downregulation. These regions retain Hop1 and continue to break in pachynema despite normal synaptonemal complex deposition. Differential retention of Hop1 requires the disassemblase Pch2/TRIP13, which preferentially removes Hop1 from telomere-distant sequences, and is modulated by the histone deacetylase Sir2 and the nucleoporin Nup2. Importantly, the uniform size of EARs among chromosomes contributes to disproportionately high DSB and repair signals on short chromosomes in pachynema, suggesting that EARs partially underlie the curiously high recombination rate of short chromosomes.

---

Meiosis generates haploid sex cells using two consecutive chromosome segregation events that follow a single cycle of DNA replication. To assist proper separation of the homologous chromosomes in the first segregation phase (meiosis I), numerous DNA double-stranded breaks (DSBs) are introduced by the topoisomerase-like enzyme Spo11 to stimulate the formation of crossover recombination products (COs). Together with sister chromatid cohesion, COs connect homologous chromosome pairs and promote their correct alignment on the meiosis I spindle ^
1-3
^.

Because DSBs are potentially genotoxic, a number of processes choreograph DSB formation at the right place and time to maintain genome integrity ^
1,3-7
^. At the chromatin level, DSB formation occurs preferentially at hotspots that depend strongly on chromatin accessibility and appropriate histone modifications ^
1,8
^. In addition, Spo11 activity is modulated over larger chromosomal domains by the specialized loop-axis architecture of meiotic chromosomes. In this architecture, DSB hotspots are primarily found on the chromatin loops, but are thought to translocate to the axial element to encounter accessory proteins necessary for DSB formation ^
9-12
^. A stimulatory role of axial-element proteins in DSB formation is supported by the fact that mutants lacking axial-element proteins exhibit severely reduced DSB levels ^
13-15
^. Indeed, the enrichment profile of axial-element proteins along chromosomes correlates well with DSB levels ^
10,11,16
^, suggesting that controlled distribution of these proteins is an important mechanism for governing the regional distribution of DSB acitivity.

In addition to the spatial regulation of Spo11 activity, a network of checkpoint mechanisms controls the timing of DSB formation ^
3,4,6,8
^. These mechanisms establish a defined window of opportunity for DSB formation by preventing DSB formation during pre-meiotic DNA replication as well as upon exit from meiotic prophase ^
17-26
^. Checkpoint mechanisms also suppress redundant DSB formation in the vicinity of already broken DNA ^
27-31
^. In addition, DSBs are progressively down-regulated as prophase proceeds. Studies suggest that the synaptonemal complex (SC), an evolutionarily conserved proteinaceous structure that assembles between homologous chromosomes, is responsible for this process ^
25,32-34
^. SC is thought to ensure cessation of DSB formation in a chromosome-autonomous fashion and likely triggers DSB downregulation following initiation of the obligatory CO on a given chromosome pair ^
25,32-36
^.

In *S. cerevisiae*, SC-dependent down-regulation of DSB activity is linked to the chromosomal reduction of the axis-associated HORMA-domain protein Hop1, which normally recruits DNA break machinery to the meiotic chromatin ^
11,33^. Reduction of Hop1 on chromosomes occurs concomitantly with SC assembly and depends on SC-mediated recruitment of the AAA^+^-ATPase Pch2 ^
33,37-39
^. In the absence of Pch2, Hop1 signal continues to accumulate on chromosome spreads in late prophase. A similar process is observed in mouse spermatocytes ^
34,40
^. Intriguingly, not all DSB hotspots in yeast are equally dampened. A number of hotspots, including several that are widely used as model hotspots (e.g. *YCR047C* and *HIS4LEU2* – a modified hotspot at *YCL030C*), remain active irrespective of the presence of the SC ^
17,33,41
^. The origin and purpose of these long-lived hotspots is not known.

One possible function of long-lived hotspots is to increase the window of opportunity for DSB formation on short chromosomes. Short chromosomes exhibit elevated recombination density in many organisms ^
25,42-46
^. In yeast, this bias is already apparent at the level of DSB formation ^
47-50
^ and is likely driven by two independent mechanisms, both of which remain poorly understood. The first mechanism causes a biased enrichment of axis proteins and DSB factors on short chromosomes and is independent of DSB formation ^
11,16
^. The second mechanism is thought to involve the SC-dependent down-regulation of DSBs linked to homologue engagement for DSB repair ^
25
^. It has been proposed that shorter chromosomes may be slower at engaging with their homologue, leading to prolonged DSB activity specifically on these chromosomes ^
25
^. Accordingly, in *zip3* mutants, which fail to implement controlled CO repair ^
51,52
^, DSB formation continues on all chromosomes, and the biased increase in DSB levels on short chromosomes is no longer detectable ^
25
^. This model, however, is likely incomplete because it predicts that long-lived hotspots will be restricted to short chromosomes. Instead, long-lived hotspots are also observed on long chromosomes ^
33
^.

Here, we show that most long-lived hotspots are located within large chromosome <u>e</u>nd-<u>a</u>djacent regions (EARs) that retain Hop1 and DSB markers in late prophase. Establishment of Hop1 enrichment in EARs requires Pch2, which preferentially removes Hop1 from interstitial chromosomal sequences, and is modulated by the histone deacetylase Sir2 and the nucleoporin Nup2. As EAR lengths are invariant between chromosomes, EARs comprise a proportionally larger fraction of short chromosomes. We propose that the spatial bias in Hop1 enrichment increases relative DSB activity on shorter chromosomes and at least in part explains the increased recombination density on short chromosomes.

## RESULTS

### Continued DSB formation is linked to chromosomal position

To identify features that distinguish short- and long-lived hotspots, we expanded the number of hotspots whose lifespan has been classified using Southern assays. To exclude dampening of DSB activity because of prophase exit ^
20
^, we deleted the *NDT80* gene, which encodes a transcription factor necessary for initiating the prophase exit program ^
53,54
^. *ndt80Δ* mutants halt meiotic progression at late prophase with fully synapsed chromosomes (pachynema) and extend the permissive time window for DSB formation ^
17,41
^, allowing more efficient capture of long-lived hotspots. Southern analysis of *ndt80Δ* cells undergoing a synchronous meiotic time course revealed new examples of long-lived (*YOL081W, YFL021W*) and short-lived hotspots (*YER004W, YER024W, YOR001W*; **
Figure 1A
**, data not shown), indicating that both hotspot classes are common in the yeast genome.

**Figure 1.**
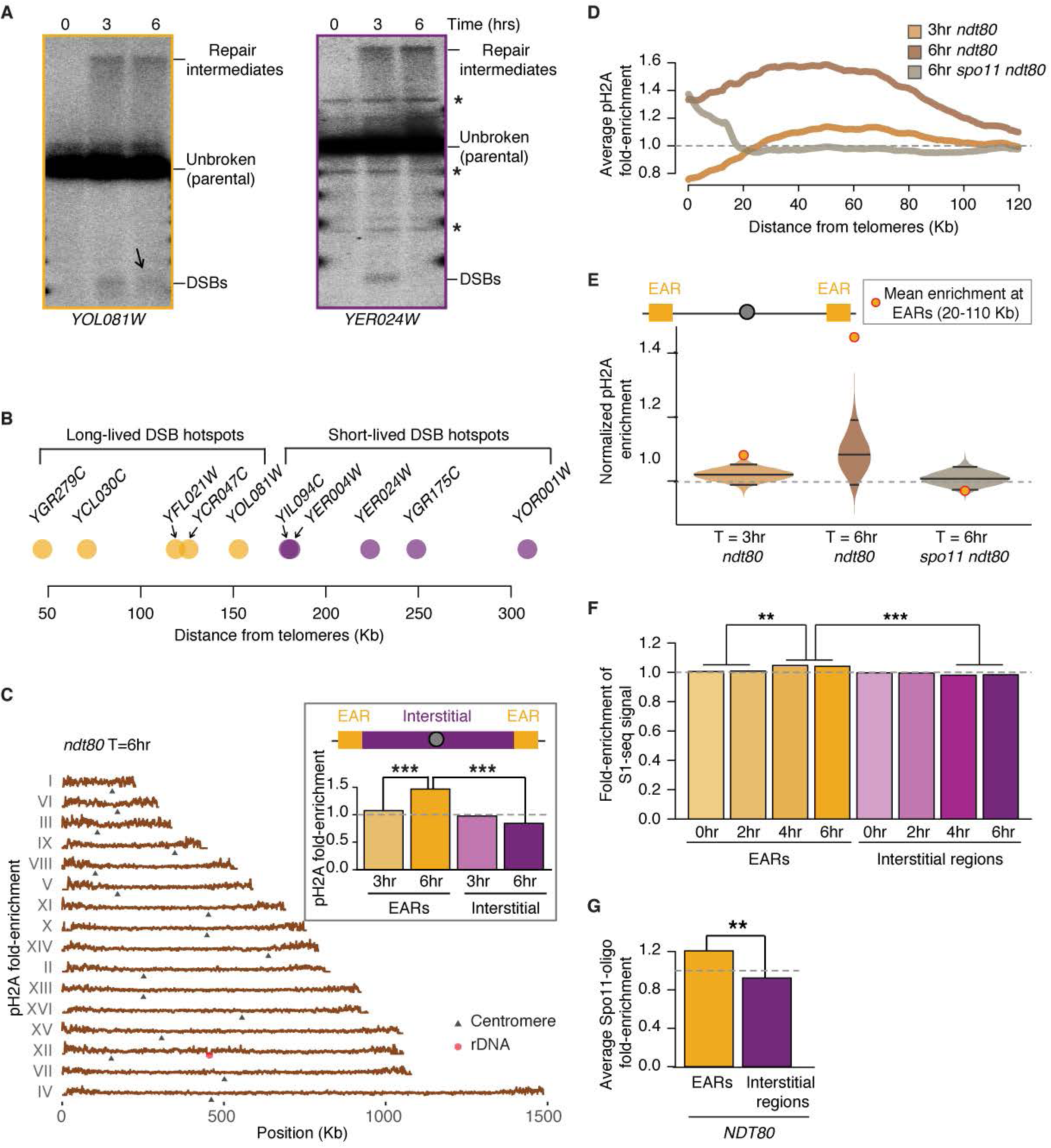
Long-lived DSB hotspots occur primarily in EARs. (A) Southern analysis to monitor DSBs from *ndt80Δ* cells progressing synchronously though meiotic prophase at *YOL081W* (a long-lived DSB hotspot, orange outline) and *YER024W* (a short-lived DSB hotspot, magenta outline). Black arrow points to continued DSBs in late prophase (T=6hrs) at the *YOL081W* DSB hotspot. * nonspecific bands. (B) Distance of a few queried DSB hotspots (orange, long-lived; magenta, short-lived) from their closest telomere (^
17,33,41
^; this manuscript, and data not shown). (C) pH2A ChIP-seq enrichment is plotted along each of the 16 yeast chromosomes, black triangles mark the centromeres and the red hexagon marks the rDNA locus. The data are normalized to a global mean of 1. Inset shows mean enrichment at EARs (20-110 Kb from telomeres; orange) and interstitial chromosomal regions (>110 Kb from telomeres; magenta) in early prophase (T=3hrs) and late prophase (T=6hrs). *** *P* < 0.001, Mann-Whitney-Wilcoxon test. (D) Mean pH2A enrichment in the EARs (32 domains) is plotted as a function of distance from telomeres. The dotted light grey line depicts genome average. (E) Bootstrap-derived distributions from ChIP-seq data are shown as violin plots. Lower and upper quantile (95% confidence intervals, CI) as well as the median computed from the bootstrap data are depicted as horizontal lines. The orange/red dot shows the mean ChIP-seq enrichment in EARs (20-110 Kb) for the respective samples. The grey dotted line is the genome average. (F) Time series of S1-seq signal reflecting resected DSB ends ^
59
^ are normalized to genome average and plotted as mean signal in EARs (20-110 Kb, 32 domains) and interstitial chromosomal regions (>110 Kb from either end of all chromosomes, 16 domains). The grey dotted line is the genome average. *** *P* < 0.001 and ** *P* < 0.01, ANOVA on mean enrichment followed by a post-hoc Tukey test. (G) Spo11-oligo within hotspots ^
25,49
^ are normalized to genome average and plotted as mean signal in EARs (20-110 Kb, 32 domains) and interstitial chromosomal regions (>110 Kb from either end of all chromosomes, 16 domains). The grey dotted line is the genome average. ** *P* < 0.01.

Plotting the positions of these and previously published hotspots analyzed in *ndt80Δ* mutants revealed that the differences in temporal regulation correlated closely with distance from telomeres. Whereas short-lived hotspots were located interstitially on chromosomes, long-lived hotspots were primarily found in large domains adjacent to chromosome ends (**
Figure 1B
**). These data suggest that continued hotspot activity is linked to chromosomal position.

To extend this analysis across the genome, we assessed markers of DSB formation by ChIP-seq assay. Histone H2A phosphorylated on serine 129 (pH2A) is a well-documented chromatin modification homologous to mammalian γ-H2AX that is activated by DSB formation and spreads into an approximately 50-Kb region around DNA breaks ^
55,56
^. Samples were collected from synchronous *ndt80Δ* cultures at time points corresponding to early prophase (T=3hrs) and late/extended prophase (T=6hrs), followed by deep sequencing of the pH2A chromatin immunoprecipitate. These analyses showed that, in early prophase, pH2A is distributed relatively evenly along chromosomes, with particular enrichment at meiotic axis sites but depletion at DSB hotspots (**Figures S1A-C**). In late prophase, however, pH2A enrichment was strongly biased towards the ends of all 16 chromosomes (**
Figure 1C
**). This enrichment was most pronounced within 20-110 Kb from telomeres (**
Figure 1D
**). We refer to these regions as chromosome <u>e</u>nd-<u>a</u>djacent regions (EARs). Averaging across all EARs revealed that this spatial bias was also apparent in early prophase, albeit to a lesser extent (**
Figures 1C
** (inset) **and 1D**). At both time points, pH2A enrichment in the EARs was above the 95% confidence interval (CI) of a bootstrap-derived distribution (**
Figure 1E
**).

pH2A enrichment near telomeres was largely dependent on *SPO11*, indicating that these regions experience enhanced meiotic DSB activity (**
Figures 1D and 1E**). Consistently, ChIP-seq analysis of Rad51, a DSB repair protein, also showed an enrichment of signal in EARs in late prophase (**Figures S1D and S1E**). We note that pH2A enrichment persisted within 20 Kb from telomeres in *spo11Δ* mutants, in line with previous observations showing DSB-independent enrichment in these regions in mitotic cells ^
57,58
^. These observations suggest that DSB activity in EARs is prolonged relative to genome average.

To assess if elevated DSB activity in EARs is also detectable in wild-type cells (*NDT80*), we analyzed publicly available genome-wide S1-seq datasets ^
59
^. S1-seq measures unrepaired DNA ends and S1 nuclease-sensitive repair intermediates and thus also reports on DSB occurrence. S1-seq signal became significantly enriched in EARs over time compared to interstitial chromosomal sequences (**
Figure 1F
**), closely mirroring the temporal enrichment of pH2A and Rad51 in these regions. This trend remained even after excluding the 3 shortest chromosomes, which consist primarily of EARs (**Figure S1F**). Analysis of available datasets ^
25,49
^ further showed that hotspot-associated Spo11-oligos, a byproduct of DSB formation, are also derived from EARs at significantly higher levels compared to telomere-distal regions (T=4hrs; **
Figure 1G and S1G**). Together, these data indicate that hotspots located in EARs are partially refractory to DSB down-regulation in late prophase.

### Domains of continued DSB formation correlate with enrichment of Hop1

We sought to identify regulators mediating the differential DSB activity in late prophase. DSB activity depends on Hop1 and correlates well with the presence of Hop1 on chromosome spreads and in genome-wide assays ^
11,14,16,33
^. Therefore, we monitored the evolution of Hop1 enrichment on wild-type (*NDT80*) meiotic chromosomes by ChIP-seq (**Figures S2A-C**). At the time of pre-meiotic DNA replication (T=2hrs), Hop1 was enriched in large domains (∼100 Kb) around the centromeres (>95% CI; **
Figures 2A, S2A and S2D**), likely linked to the early enrichment of Spo11 in these regions ^
60
^. By early prophase (T=3hrs), Hop1 enrichment became more distributed and formed peaks of enrichment along all chromosomes (**Figure S2B**), matching previously defined sites of enrichment ^
16
^. Importantly, by mid/late prophase (T=4hrs), Hop1 enrichment trended towards the EARs (**
Figures 2B and S2C**). The increase of Hop1 enrichment in EARs was even more prominent in the extended prophase of *ndt80Δ*-arrested cells (T=6hrs; **
Figures 2C and D**). In both wild type and *ndt80Δ* mutants, the increase in Hop1 enrichment was above 95% CI for a bootstrap-derived distribution of enrichment along the genome (**
Figure 2E
**). These data suggest that continued DSB formation in the EARs is the result of persistent Hop1 enrichment in these regions.

**Figure 2.**
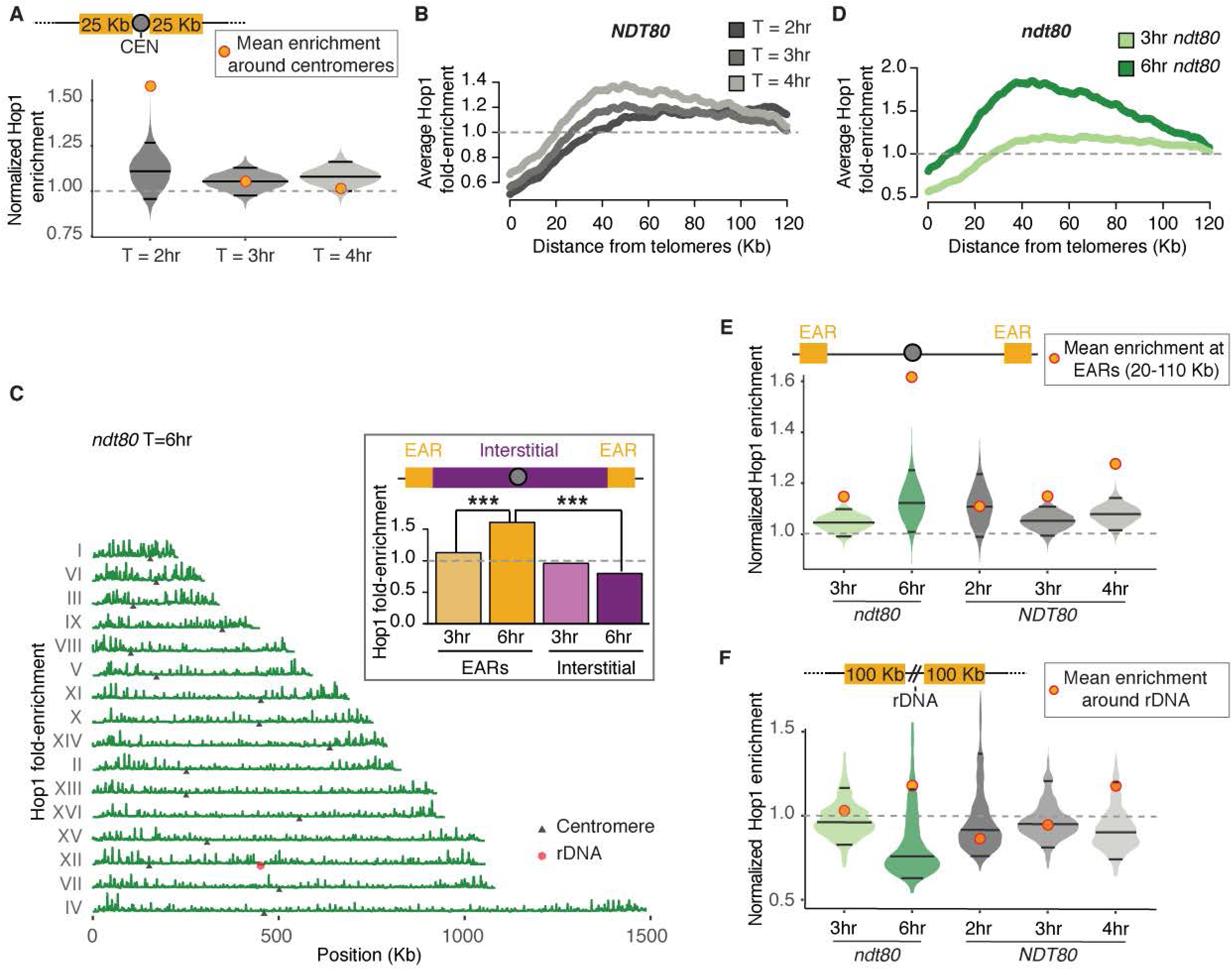
Hop1 enrichment in EARs as well as centromere and rDNA-proximal regions during prophase. (A) Time series of Hop1 enrichment around centromeres (50 Kb) in *NDT80*. Bootstrap-derived distributions are shown as violin plots and the horizontal lines within the plots represent the median and the two-ended 95% CIs. The mean Hop1 ChIP-seq enrichment around centromeres (50 Kb centered around centromeres) is shown as orange/red dots. (B) Mean Hop1 enrichment in the EARs (32 domains) is plotted as a function of distance from telomeres in *NDT80*. The grey dotted line is genome average. (C) Hop1 ChIP-seq enrichment (dark green) in *ndt80Δ*-arrested late prophase cells plotted along each of the 16 yeast chromosomes, black triangles mark the centromeres and the red hexagon marks the rDNA locus. The data are normalized to a global mean of 1. Inset shows mean enrichment in EARs (20-110 Kb, orange) and interstitial chromosomal regions (magenta) in early prophase (T=3hrs) and late prophase (T=6hrs). *** *P* < 0.001, Mann-Whitney-Wilcoxon test. (D) Mean Hop1 enrichment in the EARs (32 domains) is plotted as a function of distance from telomeres in *ndt80Δ*. (E) Bootstrap-derived distributions from Hop1 ChIP-seq data depicted as violin plots. Additionally, the horizontal lines in the violin plots represent the median and the two-ended 95% CIs. The mean Hop1 ChIP-seq enrichment in EARs (20-110 Kb) for the respective samples is shown as orange/red dots. (F) Bootstrap-derived distributions from Hop1 ChIP-seq data illustrated as violin plots. The horizontal lines in the violin plots represent the median and the two-ended CIs. The mean Hop1 ChIP-seq enrichment in rDNA-adjacent domains (100 Kb on either side of the rDNA) for the respective samples is shown as orange/red dots.

To test if the redistribution of Hop1 in late prophase reflects an overall reorganization of the meiotic chromosome axis ^
37
^, we analyzed enrichment of the chromosome axis factor Red1 by ChIP-seq (**Figure S3A**). Similar to Hop1, Red1 enrichment in EARs became more prominent in late prophase in *ndt80Δ* mutants (**Figures S3B**). Red1 enrichment in EARs was above 95% CI compared to a bootstrap-derived enrichment along the genome (**Figure S3C**) and significantly different from enrichment at telomere-distal regions (**Figure S3A**, inset). The enhanced enrichment of axis proteins in the EARs suggests that meiotic chromatin remains poised for DSB formation in these regions during late prophase.

Because Hop1 recruits Mek1 kinase to meiotic chromosomes in response to DSB-induced checkpoint activation ^
61,62
^, we also assessed Mek1 enrichment along chromosomes by ChIP-seq analysis in *ndt80Δ* cells (**Figure S3D**). Mek1 was enriched along the chromosomes in early prophase with specific enrichment at sites of axis protein binding and DSB hotspots (**Figures S4A-D**), as well as centromeres and tRNA genes (**Figures S4E and S4F**). Whereas Mek1 enrichment at axis sites persisted into late prophase, enrichment at hotspots was somewhat diminished, likely reflecting a global reduction in DNA breakage in late prophase (**Figures S4B and S4D**). Importantly, Mek1 enrichment was significantly enhanced in the EARs in late prophase (**Figures S3C-E**), providing further support that DSBs continue to form in these domains.

### EAR-like regions flanking the ribosomal DNA

In addition to the EARs, Hop1 enrichment in late prophase also increased in ∼100 Kb regions flanking the repetitive ribosomal DNA (rDNA) locus on chromosome XII (**
Figure 2C
**, red hexagon). This increase in enrichment was above the 95% CI of a bootstrap-derived distribution in *ndt80Δ* samples (**
Figure 2F
**). A similar trend, albeit below the 95% CI, was observed in wild-type (*NDT80*) cultures. Similar to EARs, the rDNA-adjacent Hop1 enrichment was accompanied by a significant local increase in pH2A signals (**Figure S3F**). Mek1 enrichment also followed this trend but was below the 95% CI. To test if pH2A enrichment reflected continued breakage of DNA in regions surrounding the rDNA, we measured DSB activity at the rDNA-adjacent *YLR152C* locus using Southern analysis. Although hotspots near the rDNA are generally weak^63^, analysis in an *ndt80Δ* background revealed that DSBs and repair intermediates increasethroughout the time course at *YLR152C* (**Figure S3G**), similar to long-lived hotspots in EARs (**
Figure 1A
**). These findings demonstrate ongoing DSB activity in late prophase next to the rDNA and suggest that the rDNA-adjacent regions, like EARs, escape negative feedback regulation of DSBs.

### The SC protein Zip1 is equally present in EARs and interstitial regions

Because cytological assays indicate that Hop1 is depleted from meiotic chromosomes upon SC assembly ^
33,37,38,64
^, we asked if EARs are less likely to assemble an SC than interstitial chromosomal sequences. To this end, we surveyed localization of the SC protein Zip1 on late-prophase chromosomes in *ndt80Δ* samples by ChIP-seq (**
Figure 3A
**). Consistent with previous reports, Zip1 was enriched around centromeres (**
Figure 3B
**). However, enrichment of Zip1 in the EARs was not different from interstitial regions (*P* = 0.564; **
Figures 3A
** (inset)**, 3C and 3D**). These findings suggest that Zip1 assembly on chromosomes is not sufficient for the spatial regulation of Hop1 in late prophase and that EARs remain enriched for Hop1 despite the presence of Zip1 in these regions.

**Figure 3.**
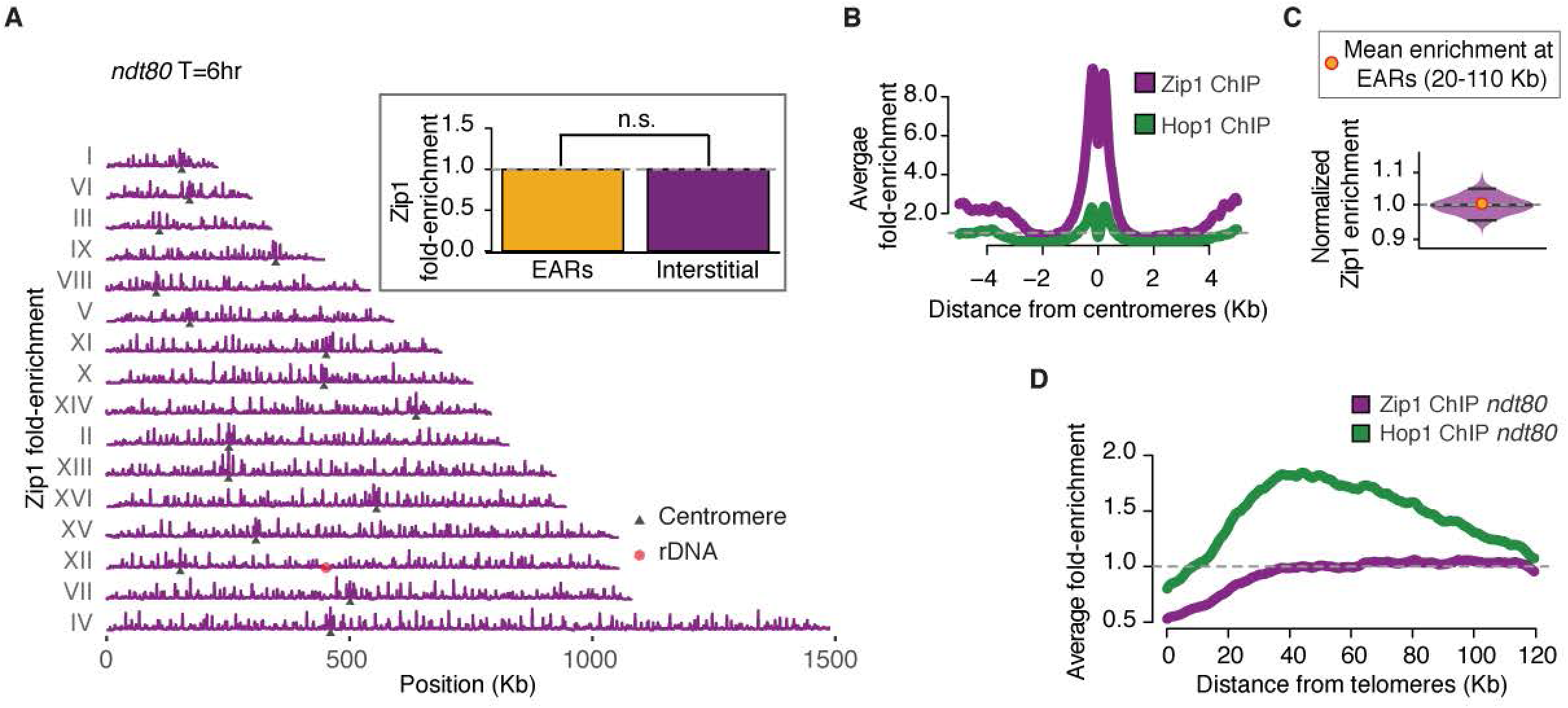
Zip1 is not depleted from EARs. (A) Zip1 ChIP-seq enrichment (magenta) in *ndt80Δ*-arrested late prophase cells plotted along each of the 16 yeast chromosomes, black triangles mark the centromeres and the red hexagon marks the rDNA locus. The data are normalized to a global mean of 1. Inset shows mean enrichment in EARs (20-110 Kb, orange) and interstitial chromosomal regions (magenta) in late prophase (T=6hrs). *P* = 0.564 (n.s.), Mann-Whitney-Wilcoxon test. (B) Zip1 (magenta) and Hop1 (dark green) ChIP-seq enrichment around centromeres from *ndt80Δ*-arrested cultures in late prophase (T=6hrs). (C) Bootstrap-derived distributions from Zip1 ChIP-seq data depicted as violin plots. The horizontal lines in the violin plots represent the median and the two-ended 95% CIs. The mean Zip1 ChIP-seq enrichment in EARs (20-110 Kb) is shown as orange/red dots.(D) Hop1 (dark green) and Zip1 (magenta) ChIP-seq enrichment in late prophase (6hrs) is plotted as a mean enrichment in the EARs (32 domains) as a function of the distance from telomeres. The grey dotted line is genome average.

### Pch2 is required for late prophase EAR enrichment of Hop1

The AAA^+^-ATPase Pch2 is recruited to synapsed chromosomes in an SC-dependent manner and is responsible for removal of Hop1 from chromosomes ^
33,37,38
^. To test if Pch2 is responsible for establishing the Hop1-enriched EARs in late prophase, we determined Hop1 binding in synchronous *pch2Δ ndt80Δ* cultures by ChIP-seq. These analyses showed abundant binding of Hop1 along chromosomes into late prophase, consistent with the persistent cytological signal of Hop1 in *pch2Δ* mutants (**
Figure 4A
**) ^
33,37,38
^. Intriguingly, the Hop1 enrichment pattern of *pch2Δ* mutants in late prophase (T=6hrs) was opposite of wild type. Hop1 specifically accumulated in interstitial regions but dropped significantly below genome average in the EARs (**
Figures 4A-C**). The altered Hop1 enrichment was reflected in altered DSB distribution and dynamics. Hotspots in interstitial regions continued to break and accumulate repair intermediates in *pch2Δ ndt80Δ* cultures, whereas hotspots in EARs exhibited comparatively reduced activity (**
Figure 4D
**). Consistently, Spo11-oligo analysis of *pch2Δ* mutants showed significantly reduced signal in EARs compared to wild type (T=4hrs; **
Figure 4E
**). These findings suggest that Pch2 promotes the removal of Hop1 from interstitial regions, leading to relative Hop1 enrichment and DSB activity in the EARs in late prophase.

**Figure 4.**
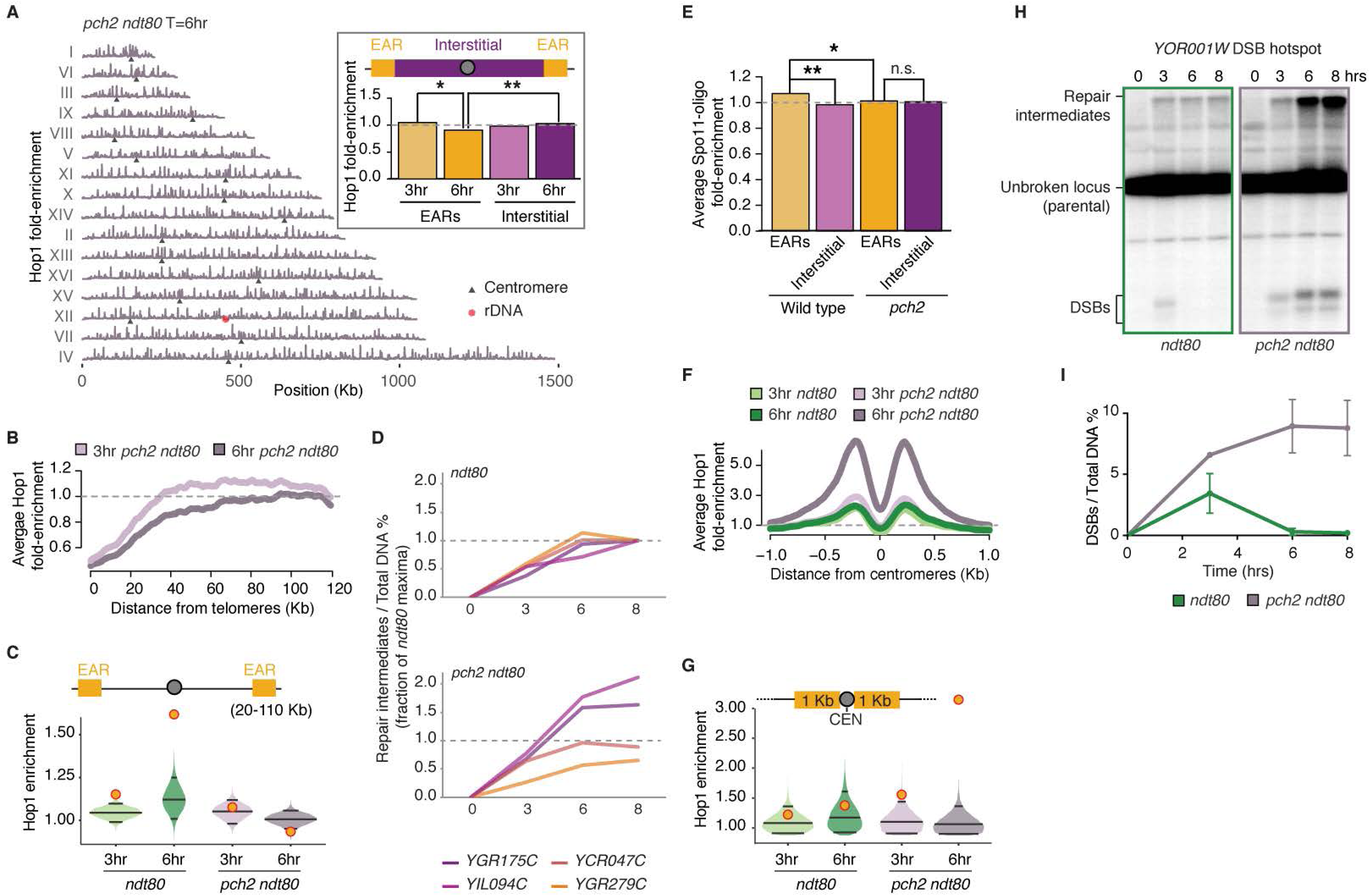
Pch2 controls regional distribution of Hop1 and DSBs in prophase. (A) Hop1 ChIP-seq enrichment in *pch2Δ ndt80Δ* late prophase (6hrs) cells plotted along each of the 16 yeast chromosomes, black triangles mark the centromeres and the red hexagon marks the rDNA locus. The data are normalized to a global mean of 1. Inset shows mean Hop1 enrichment in EARs (20-110 Kb, orange) and interstitial chromosomal regions (magenta) in early prophase (T=3hrs) and late prophase (T=6hrs). ** *P* < 0.01 and * *P* < 0.05, Mann-Whitney-Wilcoxon test. (B) Mean Hop1 enrichment in the EARs (32 domains) is plotted as a function of the distance from telomeres in *pch2Δ ndt80Δ* at early (T=3hrs) and late prophase (T=6hrs). The grey dotted line is genome average. (C) Bootstrap-derived distributions from Hop1 ChIP-seq data illustrated as violin plots for *ndt80Δ*, and *pch2Δ ndt80Δ*. The horizontal lines in the violin plots represent the median and the two-ended 95% CIs. The mean Hop1 ChIP-seq enrichment in EARs (20-110 Kb) for the respective samples is shown as orange/red dots. The grey dotted line is genome average. (D) Cumulative DNA breaks measured as percentage of repair intermediates over total DNA and depicted as a fraction of *ndt80Δ* at 8 hr timepoint are shown for *ndt80Δ*, and *pch2Δ ndt80Δ*. Interstitial hotspots (*YGR175C, YIL094C*) are shown in shades of magenta and hotspots in EARs (*YCR047C*, YGR279C) are depicted in shades of orange. (E) Spo11-oligo signals from wild type control ^
96
^ and *pch2Δ* mutant are normalized to genome average and plotted as mean signal in EARs (20-110 Kb, 32 domains) and interstitial chromosomal regions (>110 Kb from either end of all chromosomes, 16 domains). The grey dotted line is the genome average. ** *P* < 0.01, * *P* < 0.1, n.s. not significant, ANOVA on mean enrichment followed by a post-hoc Tukey test. (F) Mean Hop1 enrichment around the centromeres normalized to genome average as a function of the distance from centromeres in *ndt80Δ*, and *pch2Δ ndt80Δ*. The grey dotted line is genome average. The grey dotted line is genome average. (G) Bootstrap-derived distributions (2 Kb) of Hop1 ChIP-seq data from *ndt80Δ*, and *pch2Δ ndt80Δ* mutants are illustrated as violin plots. The horizontal lines in the violin plots represent the median and the two-ended 95% CIs. The mean Hop1 ChIP-seq enrichment around centromeres (2 Kb) for the respective samples is shown as orange/red dots.(H) Southern analysis to monitor DSBs at the *YOR001W* DSB hotspot near *CEN15* (upper panel) in *ndt80Δ*, and *pch2Δ ndt80Δ*. Percentage of DSBs over total DNA at the *YOR001W* locus at the indicated time points is shown in (I). The data are mean of two independent biological replicates and error bars represent the range.

### Pch2 suppresses Hop1 binding and DSBs at rDNA borders and centromeres

In addition to the broad effect on interstitial regions, we noted several genomic landmarks that were particularly affected by the loss of *PCH2*. As previously reported ^
63
^, Hop1 enrichment in *pch2Δ* mutants was increased around the rDNA array, resulting in elevated DSB levels in these regions (**Figure S5A**). In late prophase, Hop1 ChIP enrichment was further enhanced and DSBs continued to form near the rDNA (**Figures S5A and S5B**).

Hop1 enrichment was also strongly elevated in the immediate vicinity of centromeres in late prophase (**
Figure 4F
**). This increase was already detectable above the 95% CI in early prophase and became even more pronounced in late prophase (**
Figure 4G
**). Accordingly, Southern analysis of a centromeric DSB hotspot (*YOR001W*) revealed elevated and persistent DSB activity in *pch2Δ ndt80Δ* mutants in late prophase (**
Figures 4H and 4I**). These data indicate that Pch2 is required to restrict Hop1-linked DSB activity not only around the rDNA but also around centromeres.

### The nucleoporin Nup2 promotes Pch2 localization to chromosomes

We sought to identify additional regulators that drive Hop1 enrichment in the EARs. The disruption of several telomeric regulators, including the tethering factor Esc1, the telomere-length regulator Tel1, or the silencing factor Sir3 did not significantly affect enrichment of Hop1 in the EARs (**Figure S6A** and data not shown). Deletion of the meiotic telomere-clustering factor Ndj1 severely disrupted Hop1 enrichment in EARs, while conditional depletion caused only slight effects (**Figures S6A, S6B**). However, the interpretation of these results is complicated by the fact that loss of *NDJ1* also causes synapsis defects ^
65
^. As the nonessential nucleoporin Nup2 was recently identified as an interactor of Ndj1 ^
66
^, we also analyzed Hop1 in *nup2Δ* mutants. Intriguingly, Hop1 did not become enriched in the EARs in late prophase in *nup2Δ ndt80Δ* mutants (**
Figures 5A-C**). This effect, however, was due to diminished Hop1 removal from interstitial regions, which continued to exhibit prominent Hop1 peaks in late prophase (**
Figure 5A
**), similar to *pch2Δ* mutants. Indeed, although the phenotypes are generally less pronounced than in a *pch2Δ* mutant, loss of Nup2 also caused retention of Hop1 on synapsed chromosomal regions (**
Figures 6A and S6C**). Additionally we observed a *pch2*-like enrichment of Hop1 in the vicinity of centromeres in Hop1 ChIP-seq (**
Figure 5D
**), and increased DSB levels at interstitial hotspots by Southern assays (**Figure S6D**). These data suggest that Nup2 and Pch2 act in a common pathway.

**Figure 5.**
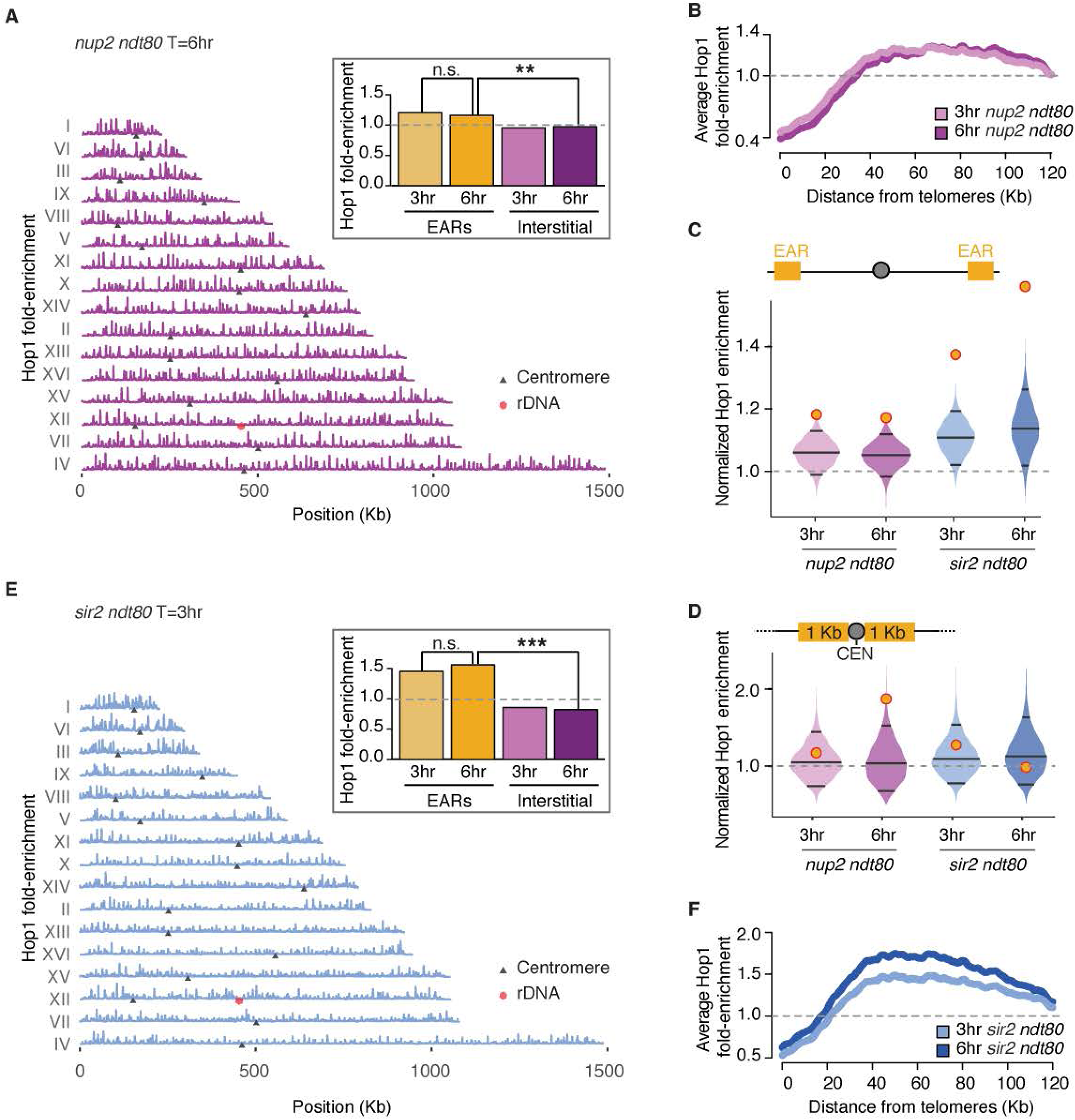
Nup2 and Sir2 also regulate regional Hop1 distribution and DSBs in prophase. (A) Hop1 ChIP-seq enrichment in late prophase cells (6hrs) from *nup2Δ ndt80Δ* in pink is plotted along each of the 16 yeast chromosomes, black triangles mark the centromeres and the red hexagon marks the rDNA locus. The data are normalized to a global mean of 1. Inset shows mean Hop1 enrichment in EARs (20-110 Kb, orange) and interstitial chromosomal regions (magenta) in early prophase (T=3hrs) and late prophase (T=6hrs). ** *P* < 0.01, and n.s. not significant, Mann-Whitney-Wilcoxon test. (B) Mean Hop1 enrichment in the EARs (32 domains) is plotted as a function of the distance from telomeres in *nup2Δ ndt80Δ*. (C) Bootstrap-derived distributions from Hop1 ChIP-seq data illustrated as violin plots for *nup2Δ ndt80Δ* (pink), and *sir2Δ ndt80Δ* (blue). The horizontal lines in the violin plots represent the median and the two-ended 95% CIs. The mean Hop1 ChIP-seq enrichment in EARs (20-110 Kb) for the respective samples is shown as orange/red dots. The grey dotted line is genome average. (D) Bootstrap-derived distributions (2 Kb) of Hop1 ChIP-seq data from *nup2Δ ndt80Δ* (pink), and *sir2Δ ndt80Δ* mutants (blue) are illustrated as violin plots. The horizontal lines in the violin plots represent the median and the two-ended 95% CIs. The mean Hop1 ChIP-seq enrichment around centromeres (2 Kb) for the respective samples is shown as orange/red dots. (E) Hop1 ChIP-seq enrichment in late prophase cells (6hrs) from *sir2Δ ndt80Δ* in blue is plotted along each of the 16 yeast chromosomes, black triangles mark the centromeres and the red hexagon marks the rDNA locus. The data are normalized to a global mean of 1. Inset shows mean Hop1 enrichment in EARs (20-110 Kb, orange) and interstitial chromosomal regions (magenta) in early prophase (T=3hrs) and late prophase (T=6hrs). *** *P* < 0.001, and n.s. not significant, Mann-Whitney-Wilcoxon test. (F) Mean Hop1 enrichment in the EARs (32 domains) is plotted as a function of the distance from telomeres in *sir2Δ ndt80Δ*.

**Figure 6.**
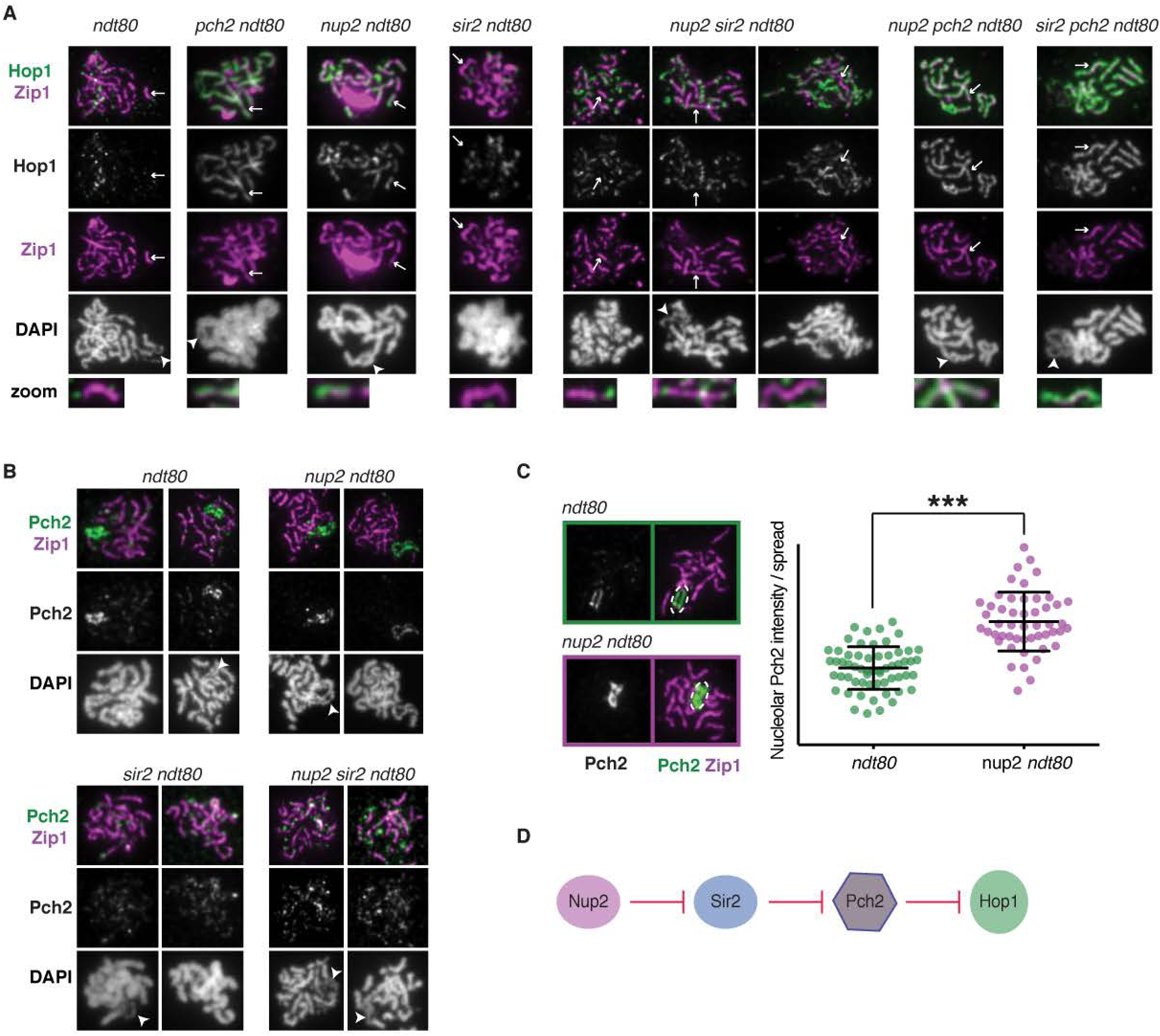
Network of Nup2, Sir2 and Pch2 regulates Hop1 on meiotic chromosomes. (A) Immunofluorescence of Hop1 (green/ grey), Zip1 (magenta), and DAPI (grey) on spread chromosomes from *ndt80Δ, pch2Δ ndt80Δ, nup2Δ ndt80Δ, sir2Δ ndt80Δ, nup2Δ sir2Δ ndt80Δ, nup2Δ pch2Δ ndt80Δ*, and *sir2Δ pch2Δ ndt80Δ* meiotic cultures. Arrow points to a synapsed region (magenta) and zoomed-in signal are shown in the last panel. In *pch2Δ ndt80Δ, nup2Δ ndt80Δ, nup2Δ pch2Δ ndt80Δ*, and *sir2Δ pch2Δ ndt80Δ* mutants, the synapsed region overlaps with Hop1 (green) but not in *sir2Δ ndt80Δ*, and *nup2Δ sir2Δ ndt80Δ* mutant samples. Arrowhead points to rDNA where easily distinguishable. (B) Immunofluorescence of Pch2 (green/ grey), Zip1 (magenta), and DAPI (grey) on spread chromosomes from *ndt80Δ, nup2Δ ndt80Δ, sir2Δ ndt80Δ*, and *nup2Δ sir2Δ ndt80Δ* meiotic cultures. Arrowhead points to rDNA where distinguishable. (C) Right Panel: Quantification of nucleolar Pch2 intensity per spread nucleus in *ndt80Δ* (green dots), and *nup2Δ ndt80Δ* (purple dots). n > 50; error bars are S.D. from the mean. Left Panel: Representative immunofluorescence on a spread nucleus for Pch2 (green) and Zip1 (magenta) in *ndt80Δ* (top), and *nup2Δ ndt80Δ* (bottom). Measured area is outlined by dotted white ovals. (D) Schematic representation of genetic interaction between Sir2, Nup2 and Pch2 for evicting Hop1 from meiotic chromosomes.

A common pathway is also supported by the fact that the defects in Hop1 localization of the single mutants are non-additive. The *nup2Δ pch2Δ* double mutant resembles the *pch2Δ* single mutant when Hop1 accumulation is analyzed on chromosome spreads (**
Figures 6A and S6C**). The fact that the double mutant phenocopies the stronger *pch2Δ* phenotype also implies that Pch2 acts downstream of Nup2. Consistent with this interpretation, Pch2 foci on chromosome spreads are noticeably diminished in the absence of Nup2, with most of the Pch2 staining being concentrated in a bright nucleolar signal (**
Figure 6B
**). Quantification of the nucleolar signal indicated that Pch2 is even more abundant in the nucleolus in *nup2Δ* mutants than in wild type (**
Figure 6C
**).

To investigate if the nucleolar pool of Pch2 is functional in the absence of Nup2, we analyzed DSB formation and Hop1 enrichment near the rDNA. *nup2Δ* mutants do not phenocopy *pch2Δ* mutants for rDNA-associated phenotypes, as there is no DSB induction near the rDNA (**Figure S5A**). In fact, *nup2Δ* mutants showed a relative decrease in Hop1 ChIP-seq signal near the rDNA compared to wild type (**Figure S5B**). These results imply that the nucleolar pool of Pch2 is fully functional in the absence of Nup2 and suggest that Nup2 is not a general activator of Pch2 function but rather acts through controling the relative nuclear distribution of Pch2.

### Nup2 regulation of Pch2 is mediated by Sir2

Nup2 may either promote the binding of Pch2 to synapsed chromosomes or suppress sequestration of Pch2 in the nucleolus. To distinguish between these possibilities, we analyzed mutants lacking the rDNA-enriched silencing factor Sir2, which is required for the nucleolar localization of Pch2 ^
38
^. *nup2Δ sir2Δ* double mutants showed a complete loss of Pch2 from the nucleolus similar to the *sir2Δ* single mutant (**
Figure 6B
**, lower panels), indicating that the nucleolar over-enrichment of Pch2 in *nup2Δ* mutants is fully dependent on Sir2. Importantly, Pch2 signal on synapsed chromosomes was indistinguishable between *sir2Δ* and *sir2Δ nup2Δ* double mutants (**
Figures 6B and S6E**), showing that Nup2 does not promote binding of Pch2 to synapsed chromosomes. These data suggest that Nup2 counteracts Sir2-dependent recruitment of Pch2 to the nucleolus.

Consistent with this model, the aberrant Hop1 accumulation on synapsed chromosomes observed in *nup2Δ* mutants is rescued in the *sir2Δ nup2Δ* double mutant. Although the *sir2Δ nup2Δ* double mutant has substantial synapsis defects, Hop1 is never observed on synapsed chromosome fragments (**
Figure 6A
**), suggesting efficient Hop1 removal by Pch2. These data are consistent with the abundant presence of Pch2 on chromosomes in the double mutant and indicate that Nup2 acts upstream of Sir2 in the control of Hop1.

To further investigate the role of Sir2 in controlling Hop1 distribution, we performed ChIP-seq of Hop1. This analysis showed that Hop1 was precociously enriched in the EARs in *sir2Δ ndt80Δ* mutants, with strong enrichment already detectable in early prophase (T=3hrs) (**
Figures 5C and 5E**). This pattern is consistent with the increased abundance of Pch2 on synapsed chromosomes and may reflect a faster progression of meiotic events when Pch2 is overactive^37^. Consistent, with an accelerated prophase, we noted a slightly faster appearance of fully synapsed chromosomes in *sir2Δ* mutants (**Figure S6F**) as well as faster accumulation of repair intermediates, in particular for interstitial regions, which lost Hop1 binding (**Figure S6D**). This acceleration likely reflects the premature elimination of Mek1 which normally suppresses repair ^33,67^. It is possible that this faster progression also leads to a premature shutdown of DSB formation because Spo11-oligo analysis at T=4hrs shows comparable levels of DSB formation in EARs and interstitial regions in *sir2Δ* mutant. (**Figure S6G**). In line with this possibility, repair intermediates do not continue to accumulate at later time points in *sir2Δ ndt80Δ* mutants (**Figure S6D**). Taken together, our data implicate a regulatory pathway consisting of Nup2, Sir2 and Pch2 in driving the enrichment of Hop1 in the EARs and controlling the window of opportunity for DSB formation (**
Figure 6D
**).

### Enrichment of Hop1 and DSB markers exhibits a size bias favoring short chromosomes

The fact that EARs occupy a proportionally much larger fraction of short chromosomes (**
Figure 7A
**) provides a possible mechanism for increasing relative DSB levels on these short chromosomes. Indeed, plotting pH2A enrichment/Kb as a function of chromosome size revealed a distinct, *SPO11*-dependent over-enrichment of pH2A on short chromosomes in late prophase (**
Figure 7B
**). A similarly biased enrichment on short chromosomes was also observed for Hop1 and Mek1 in early prophase and increased further in late prophase (**
Figure 7C and S7A**). Moreover, increasing enrichment of Hop1 on short chromosomes occurred during prophase regardless of whether *NDT80* was present (**Figure S7B**).

The early enrichment of Hop1 and Mek1 on short chromosomes may be driven by the underlying enrichment of Red1 protein, which recruits Hop1 to chromosomes ^
64,68
^ and also exhibits chromosome size bias for enrichment in early prophase (**Figure S7C**) ^
16
^. Supporting this model, the pattern of chromosome size bias between Red1 and Hop1 in early prophasewas not significantly different (ANOVA, p=0.167). However, the Red1 chromosome size bias did not increase between early and late prophase (**Figure S7C**), indicating that the late prophase enrichment of Hop1 (and Mek1) on short chromosomes occurs by a different mechanism. Importantly, calculating the mean Hop1 enrichment per chromosome while excluding EARs significantly reduces the chromosome size bias in late prophase in both *ndt80Δ*-arrested and wild-type cells (**
Figure 7D and S7D**). These analyses support the model that EARs are responsible for the late prophase enrichment of Hop1 on short chromosomes.

**Figure 7.**
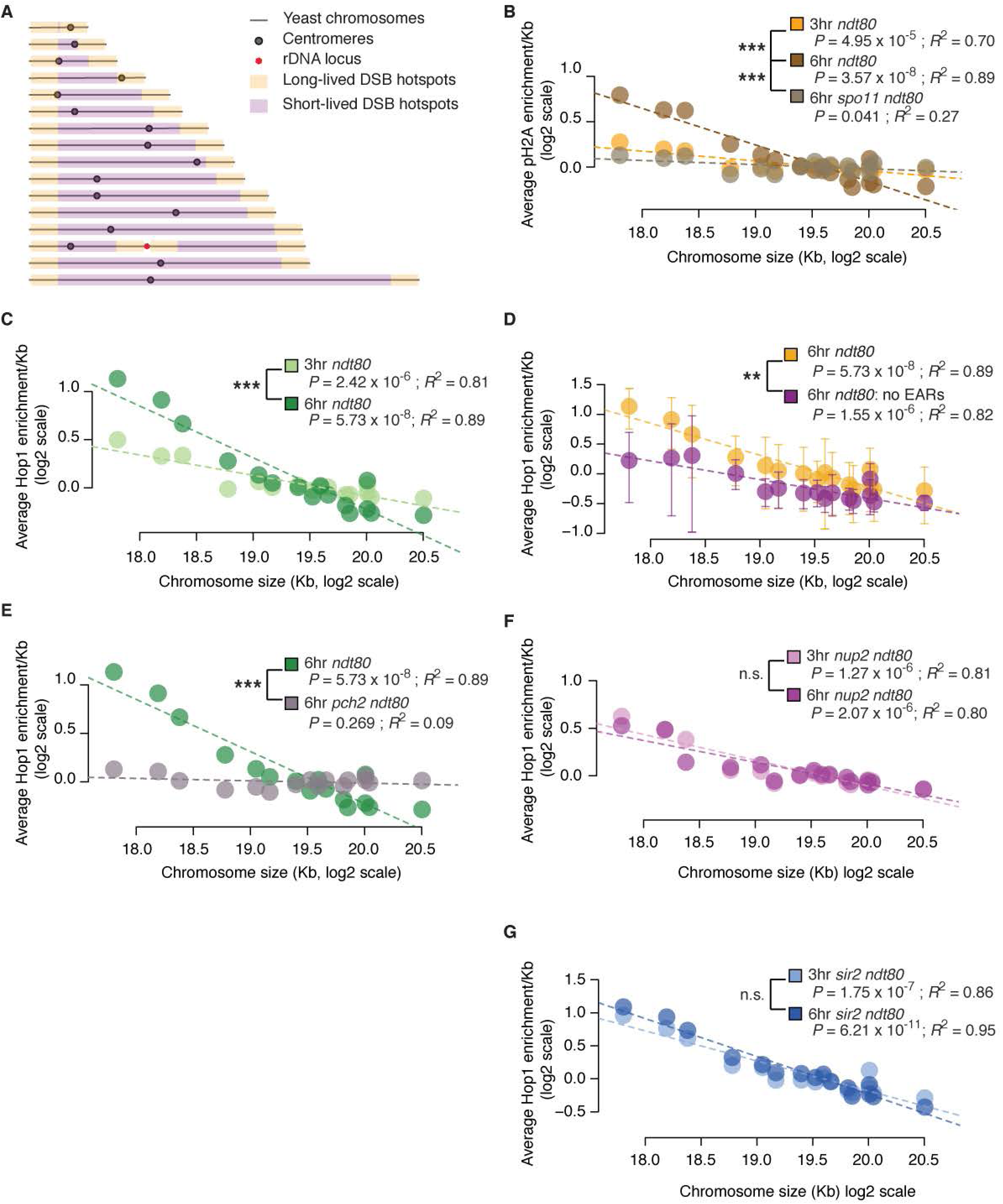
Enrichment of Hop1 and DSB markers on short chromosomes increases significantly in late prophase and depends on Pch2. (A) Schematic of telomere-adjacent enrichment of Hop1 in late prophase predicts bias for DSBs on short chromosomes. The orange bars illustrate the large regions (∼100 Kb) of long-lived DSB hotspots at telomere-adjacent and rDNA-adjacent domains. Interstitial regions (magenta) harbor mainly the short-lived DSB hotspots. (B-G) Mean ChIP-seq enrichment per Kb is plotted for each chromosome on log scale with regression analysis. *P* and *R*
^
2
^ values are noted below the sample name. *R*
^
2
^, measure of the fit of the points to the line, can vary from 0-1.0 with 1.0 indicating a perfect fit. *P* is the probability of obtaining large *R*
^
2
^ values. Two-way ANOVA was performed to test significant difference in the slope between the regression lines for different ChIP-seq samples. ANOVA-derived *P* values are indicated, *** *P* < 0.001. pH2A ChIP-seq enrichment in early (T=3hrs) and late prophase (T=6hrs) in *ndt80Δ* and late prophase (6hrs) in *spo11Δ ndt80Δ* in (B). Hop1 ChIP-seq in early (T=3hrs) and late prophase (T=6hrs) in *ndt80Δ* samples (C). Hop1 ChIP-seq in late prophase from *ndt80Δ* samples (T=6hrs) is plotted in orange (D). Plot in magenta is Hop1 enrichment from telomere-distal regions lacking EARs (110 Kb from each chromosome end). Error bars are standard deviation of the means of 10 equal sized bins for each chromosome. Late prophase (T=6hrs) enrichment of Hop1 in *ndt80Δ* and *pch2Δ ndt80Δ* cultures (E). Hop1 ChIP-seq in early (3hrs) and late prophase (T=6hrs) in *nup2Δ ndt80Δ* samples (F). Hop1 ChIP-seq in early (3hrs) and late prophase (T=6hrs) in *sir2Δ ndt80Δ*samples (G).

If EARs drive biased enrichment of Hop1 on short chromosomes then this effect should be abrogated in *pch2Δ* mutants, which do not exhibit Hop1 enrichment in EARs. Indeed, whereas the pattern of Hop1 chromosome size bias was not significantly different between *pch2Δ* and *PCH2* samples in early prophase (*P* = 0.691) (**Figure S7E**), in late prophase all chromosomes in *pch2Δ* mutants had a similar Hop1 enrichment/Kb irrespective of chromosome size (**
Figure 7E
**). Hop1 chromosome size bias was also diminished in *nup2Δ* mutants (**
Figure 7F
**) but enhanced in *sir2Δ* mutants (**
Figure 7G
**). These observations indicate that a regulatory mechanism comprised of Nup2, Sir2, and Pch2 is responsible for maintaining the chromosome-size bias in meiotic prophase.

## DISCUSSION

Our findings uncover striking regional control of DSB potential during meiosis. DSB hotspots in large domains (∼100 Kb) adjacent to chromosome ends, as well as regions bordering the rDNA locus, continue to break well after the SC down-regulates hotspots in interstitial chromosomal regions. This positional regulation increases the break potential on short chromosomes in the course of prophase and reveals an intuitive mechanism for promoting formation of the obligatory crossover on short chromosomes without having to measure chromosome length.

### Role of the SC in down-regulating hotspot activity

Research in several organisms, including yeast and mice, has strongly implicated the SC in mediating DSB down-regulation through the removal of HORMAD family proteins ^
25,33,34
^. The results reported here are fully consistent with this view but show that, at least in yeast, chromosomes have specialized domains (EARs) that escape this down-regulation. Intriguingly, EARs retain Hop1 despite normal accumulation of the SC protein Zip1. Thus, although chromosome synapsis is spatially correlated with the removal of HORMAD family proteins ^33,34,37,38^, Zip1 deposition in itself is clearly not sufficient for Hop1 eviction. In *C. elegans*, which also utilizes the SC to down-regulate DSB formation in a timely manner, CO designation is thought to lead to structural changes within the SC that prevent further DSB formation ^
69-72
^. Similar structural changes may also occur in the yeast SC and may be required for Hop1 removal ^
33,73
^. If so, then Hop1 in EARs is likely protected from these effects, either because Pch2-dependent removal is suppressed in the EARs or because Hop1 continues to load in these regions. The unique nature of EARs in this respect is highlighted by the fact that deletion of *PCH2* leads to a build-up of Hop1 only in the interstitial regions, while EARs become comparatively under-enriched. These data suggest that Hop1 binding in EARs is controlled by mechanisms that are distinct from the rest of the genome.

The molecular features that distinguish EARs from interstitial chromosomal regions remain to be discovered, although the consistent distance of EARs from chromosome ends implies a role for telomere-associated processes or the nuclear periphery. The latter possibility is particularly appealing because EAR-like domains of Hop1 enrichment are also observed near the rDNA, which is located near the nuclear envelope ^
74,75
^. However, analysis of a limited set of telomeric regulators (*TEL1, SIR3, NDJ1*) and nuclear envelope factors (*ESC1, NUP2*) did not yield regulators of EAR establishment. These results obviously do not exclude redundant mechanisms or a role for different telomeric or nuclear-envelope factors. However, it is equally possible that other dynamic chromatin features, such as differences in replication timing, gene activity, or chromatin topology govern the observed patterns of Hop1 enrichment and provide the architectural basis of EARs.

### Control of DSBs near centromeres

In addition to the broad changes in enrichment of Hop1 across chromosomes during meiotic prophase, we also observed unexpected dynamics of Hop1 around centromeres. Most notably, we found a strong centromeric enrichment of Hop1 in the earliest stages of prophase, before Hop1 has fully accumulated on chromosome arms. Centromeric Hop1 enrichment may similarly reflect nuclear architecture because prior to the tethering of telomeres to the nuclear envelope, the centromeres are clustered at the spindle pole body embedded in the nuclear envelope ^
76,77
^. Interestingly, the early prophase dynamics of Hop1 mirrors the distribution of Spo11, which is also enriched near centromeres before distributing to the arms ^
60
^. Indeed, Spo11 is likely active in these regions because we observe centromeric DSBs in early prophase. Curiously, we also detected an enrichment of Mek1 around centromeres. Recruitment of Mek1 is unexpected because Mek1 suppresses repair with the sister chromatid ^
33,62,78
^, yet DSBs at centromeres are thought to be channeled by Zip1 to primarily use the sister for recombination to protect against chromosome missegregation ^
79,80
^. Perhaps, in addition to mediating Mek1 removal from synapsed chromosomes ^
33
^, Zip1 also suppresses Mek1 activity at the centromeres without evicting it. Another unexpected finding is that Pch2 suppresses DSBs around centromeres in late prophase. However, as COs are not enhanced around the centromeres in *pch2Δ* mutants^43^, Zip1 activity must be sufficient to prevent any deleterious inter-homologue COs in this situation. These findings indicate that several mechanistic layers restrict DSBs and COs at the centromeres, highlighting the importance of limiting COs in this region.

### Regulation of Pch2 by Nup2

Our findings also offer new insights into the regulation of Pch2. We show that the nucleoporin Nup2 promotes the binding of Pch2 to synapsed chromosomes and provide genetic evidence that Nup2 functions by counteracting the histone deacetylase Sir2. This regulation may not be direct as Nup2 is primarily localized to the nuclear pores, whereas Sir2 and Pch2 are strongly enriched in the nucleolus. We note, however, that Sir2 is also present at euchromatic replication origins ^
81
^, which may also be sites of Pch2 activity ^
63
^. Furthermore, Nup2 is a mobile nucleoporin that interacts with chromatin to regulate transcription and contribute to boundary activity ^
82-85
^, which may allow for interactions with Sir2 and the regulation of Hop1 on chromosomes.

### EARs: an unbiased mechanism that contributes to bias

Our analysis of Hop1 dynamics sheds important light on the mechanistic basis of the meiotic chromosome-size bias for recombination. In several organisms, including humans, short chromosomes exhibit higher levels of recombination ^
42-46,86
^, a bias that in yeast is already apparent from elevated levels of axis protein deposition and DSB formation on short chromosomes ^
25,47-50
^. As EAR length is invariant regardless of chromosome size, EARs comprise a proportionally much larger fraction of short chromosomes (**
Figure 7A
**). The resulting bias in Hop1 enrichment could thus partially mediate the establishment of chromosome size bias in DSBs and COs.

Available data suggests that chromosome size bias for Hop1 enrichment in late prophase is a direct consequence of preferential SC-dependent removal of Hop1 from the interstitial chromosomal regions. Consistent with this notion, disrupting either CO-associated SC assembly (by deleting the CO-implementing factor *ZIP3*) or preventing the SC from removing Hop1 (by deleting *PCH2*) leads to a loss of chromosome size bias for recombination ^
25,43,87
^. Intriguingly, in both situations, the failure to remove Hop1 differentially affects the EARs. In *zip3Δ* mutants, DSB enrichment in EARs is diminished compared to wild type (*P* = 0.58, Mann-Whitney-Wilcoxon test; **Figure S7F**). Similarly, the percentage of COs and noncrossovers (NCOs) per meiosis drops significantly in the EARs of *pch2Δ* mutants, while average CO (and NCO) counts per chromosome surge with increasing chromosome size ^
43
^. We note that although short chromosomes in yeast are slowest to synapse ^
88
^, we find no evidence of reduced Zip1 accumulation on short chromosomes (**Figure S7G**). These data suggest that synaptic delays are too small to be detected by ChIP-seq analysis and therefore cannot explain the elevated Hop1 levels on short chromosomes observed by the same assay.

Intriguingly, COs are enriched in sub-telomeric regions in several organisms ^
42,89-95
^. Moreover, an ancient telomeric fusion that gave rise to human chromosome 2 led to a decrease in crossovers rates near the fused chromosome ends compared to chimpanzees, which maintained the two separate chromosomes ^
89
^. Thus, some fundamental features of EARs may well be evolutionarily conserved.

We propose that EARs provide a safety mechanism that ensures that DSB formation is not prematurely inactivated by the formation of the SC. Premature down-regulation of the DSB machinery is particularly problematic for short chromosomes because of their inherently lower number of DSB hotspots. By establishing privileged regions that are refractory to this down-regulation, cells may ensure that all chromosomes retain a (limited) potential for DSB formation and successful crossover recombination throughout meiotic prophase.

## AUTHOR CONTRIBUTIONS

Conceptualization, V.V.S. and A.H.; Investigation, V.V.S., X.Z., S.K., and A.H.; Software, V.V.S., T.E.M., L.A.V.S., and A.H.; Formal Analysis, V.V.S. and A.H.; Resources, V.V.S., P.A.S, N.M.H., and A.H.; Writing – Original Draft, V.V.S. and A.H.; Writing – Review & Editing, V.V.S., X.Z., T.E.M., L.A.V.S., P.A.S, N.M.H., S.K., and A.H.

## ACKNOWLEDGEMENTS

We are grateful to Brian Parker at NYU for advice on statistics. We thank the NYU Genomics Core facility for technical assistance and data processing. This work was funded in part by grant R01 GM111715 from the NIH and research grant #6-FY16-208 from the March of Dimes Foundation to A.H.; grant R35 GM118092 from the NIH to S.K and MSKCC Cancer Center Core Grant P30 CA008748; grant R01 GM050717 from the NIH to N.M.H.; grants BFU2015-65417-R from MINECO and CSI084U16 from Junta de Castilla y León in Spain to P.A.S.

## METHODS

### Contact for reagent and resource sharing

Further information and requests for resources and reagents should be directed to and will be fulfilled by the Lead Contact, Andreas Hochwagen.

### Experimental model and subject details

All strains used in this study are in the SK1 background and listed in the **Table S1**.

### Method details

#### Growth conditions

Synchronous meiotic time-courses were performed as described in ^
33
^. The strains were first patched on glycerol media (YPG) and then transferred to rich media with 4% dextrose (YPD 4%). The cells were then grown at 23°C for 24 hrs in liquid YPD and diluted into pre-sporulation media (BYTA) at A_600_ 0.3. The BYTA culture was grown at 30°C for 16 hrs. The cells were washed twice in sterile water and transferred to sporulation media (0.3% potassium acetate) at 30°C to induce synchronous sporulation. Samples for ChIP-seq (25 mL) or DSB Southern assays (10mL) were collected at the indicated time-points. Growth conditions for obtaining Spo11-oligo sequences are described in ^
96
^.

#### DSB Southern analysis

Meiotic cells collected at the indicated time points were embedded in agarose plugs to minimize background from random shearing and genomic DNA was extracted ^
63
^. The plugs were washed 4x 1hr in TE followed by 4x 1hr washes in the appropriate NEB buffer. Plugs for each time-point were transferred to separate tubes and melted at 65°C. The genomic DNA in molten agarose was equilibrated at 42°C prior to incubation with appropriate restriction enzyme(s). The digested DNA was electrophoresed in 0.8% agarose (Seakem LE) in 1X TBE at 80V for 18 hrs. The DNA was transferred to Hybond-XL nylon membrane (GE Healthcare) by capillary transfer and detected by Southern hybridization as described in ^
33
^. Restriction enzymes used for DSB analysis and primer sequences to construct probes are listed in the **Table S2**. Probes labeled with ^
32
^P dCTP were generated using the listed primers and a Prime-It random labeling kit (Agilent). Southern blots were exposed on an Fuji imaging screen and the phosphor-signal was detected on Typhoon FLA 9000 (GE) and quantified using ImageJ software (http://imagej.nih.gov/ij/). Plots were generated using the Graphpad program in Prism.

#### Chromatin immunoprecipitation and Illumina sequencing

Samples were collected from sporulation cultures at the indicated time points and crosslinked in 1% formaldehyde (Sigma) for 30 min. The formaldehyde was quenched with 125mM glycine. ChIP was performed as described in ^
97
^ using antibodies listed in the **Table S3**. Libraries for ChIP sequencing were prepared by PCR amplification with TruSeq adaptors (Illumina) as described in ^
16
^. Quality of the libraries was checked on 2100 Bioanalyzer or 2200 Tapestation. Libraries were quantified using qPCR prior to pooling. The ChIP libraries were sequenced on Illumina HiSeq 2500 or NextSeq 500 instruments at NYU Biology Genomics core to yield 51/50 bp single-end reads.

#### Processing of reads from Illumina sequencing

Illumina output reads were processed as described in ^
98
^. The reads were mapped to SK1 genome ^
99
^ using Bowtie ^
100
^. Only reads that mapped to a single position and also matched perfectly to the SK1 genome were retrieved for further analysis. 3’ ends of the reads were extended to a final length of 200bp using MACS2 2.1.1 (https://github.com/taoliu/MACS) and probabilistically determined PCR duplicates were removed. The input and ChIP pileups were SPMR-normalized (signal per million reads) and fold-enrichment of ChIP over input data was used for further analyses. The pipeline used to process Illumina reads can be found at (https://github.com/hochwagenlab/ChIPseq_functions/tree/master/ChIPseq_Pipeline_v3/).

#### Spo11-oligo mapping

Spo11-oligos immunoprecipitated from synchronous meiotic cultures (T=4hrs samples) as described using anti-FLAG antibody ^
96
^. The Spo11-oligos were sequenced on Illumina HiSeq in the Memorial Sloan Kettering Cancer Center (MSKCC) Integrated Genomics Operation core facility. The adaptors were clipped followed by alignment of the oligo reads to S288c (sacCer2) reference genome using a custom pipeline ^
25,50
^. Averaged maps from biological replicates were used for further analysis. Oligos within the rDNA (coordinates 451,000 and 471,000 on ChrXII) are highly enriched in *sir2Δ* datasets and were removed from all datasets prior to analysis.

### Quantification and statistical analyses

ChIP-seq data from two biological replicates were merged prior to analyses using the ChIPseq_Pipeline_v3 except for Hop1 ChIP from *sir3Δ ndt80Δ, tel1Δ ndt80Δ, ndj1Δ ndt80Δ* samples in Figure S5A and No tag *ndt80Δ*, and *NDJ1-FRB ndt80Δ* samples for Ndj1 depletion studies in Figure S5B. ChIP-seq data was normalized to global mean of one and regional enrichment was calculated. The R functions used can be found at (https://github.com/hochwagenlab/hwglabr2/).

Statistical significance tests were performed in R 3.3.3. Either Mann-Whitney-Wilcoxon test on mean enrichment by chromosome (16 chromosomes; EARs versus interstitial) was used to test for significance or one-way ANOVA with post-hoc Tukey test was used. Two-way ANOVA for multiple linear regressions with interaction was performed on log2-scaled ChIP-seq enrichment to test variation in the slopes (chromosome size bias) of two different samples. For bootstrap analyses, random samplings of the ChIP data were performed on each of the 16 circularized chromosomes and this was repeated 5000 times. The samplings were equivalent to the experimental query in size and number for each experiment. For instance, bootstrap samplings for EARs were two unlinked samplings from each of the 16 chromosomes, for the centromeres bootstrap involved only a single sampling of each of the 16 chromosmes while for the rDNA bootstrap the entire genome was sampled twice. Additionally, to assay enrichment at the rDNA borders, EARs (120 Kb from either telomere) were excluded from the genome for random bootstrap samplings. Both averaged random sampling data and experimental query were normalized to genome average. The median and two-sided 95% CI was calculated based on the spread of the bootstrap-derived distribution of enrichment.

### Data and software availability

All datasets reported in this paper (except published datasets) are available at the Gene Expression Omnibus (GEO) with the accession number GSE105111.

#### Genome-wide DSB and S1-seq datasets

Genome-wide S1-seq datasets for wild-type meiosis, GEO accession number GSE85253, were obtained from ^
59
^. The processed dataset aligned to the S288c reference genome (sacCer2) was used. The mapped Spo11-oligo counts within hotspots for wild type and *zip3Δ* mutant, GEO number GSE48299, aligned to the S288c reference genome (sacCer2) were obtained from ^
25
^. Additional wild-type Spo11-oligo counts data within hotspots, GEO number GSE71930, also aligned to the S288c reference genome (sacCer2) were obtained from ^
49
^. Wild type Spo11-Flag oligo sequencing data are from GEO dataset GSE67910 ^
96
^.

## REFERENCES

1. de Massy, B. Initiation of meiotic recombination: how and where? Conservation and specificities among eukaryotes. Annu Rev Genet 47, 563–99 (2013).

2. Hunter, N. Meiotic Recombination: The Essence of Heredity. Cold Spring Harb Perspect Biol 7, a016618 (2015).

3. Lam, I. & Keeney, S. Mechanism and regulation of meiotic recombination initiation. Cold Spring Harb Perspect Biol 7, a016634 (2014).

4. Cooper, T.J., Garcia, V. & Neale, M.J. Meiotic DSB patterning: A multifaceted process. Cell Cycle 15, 13–21 (2016).

5. Gray, S. & Cohen, P.E. Control of Meiotic Crossovers: From Double-Strand Break Formation to Designation. Annu Rev Genet 50, 175–210 (2016).

6. Subramanian, V.V. & Hochwagen, A. The meiotic checkpoint network: step-by-step through meiotic prophase. Cold Spring Harb Perspect Biol 6, a016675 (2014).

7. Yu, Z., Kim, Y. & Dernburg, A.F. Meiotic recombination and the crossover assurance checkpoint in Caenorhabditis elegans. Semin Cell Dev Biol 54, 106–16 (2016).

8. Keeney, S., Lange, J. & Mohibullah, N. Self-organization of meiotic recombination initiation: general principles and molecular pathways. Annu Rev Genet 48, 187–214 (2014).

9. Acquaviva, L. et al. The COMPASS subunit Spp1 links histone methylation to initiation of meiotic recombination. Science 339, 215–8 (2013).

10. Blat, Y., Protacio, R.U., Hunter, N. & Kleckner, N. Physical and functional interactions among basic chromosome organizational features govern early steps of meiotic chiasma formation. Cell 111, 791–802 (2002).

11. Panizza, S. et al. Spo11-accessory proteins link double-strand break sites to the chromosome axis in early meiotic recombination. Cell 146, 372–83 (2011).

12. Sommermeyer, V., Beneut, C., Chaplais, E., Serrentino, M.E. & Borde, V. Spp1, a member of the Set1 Complex, promotes meiotic DSB formation in promoters by tethering histone H3K4 methylation sites to chromosome axes. Mol Cell 49, 43–54 (2013).

13. Kim, K.P. et al. Sister cohesion and structural axis components mediate homolog bias of meiotic recombination. Cell 143, 924–37 (2010).

14. Mao-Draayer, Y., Galbraith, A.M., Pittman, D.L., Cool, M. & Malone, R.E. Analysis of meiotic recombination pathways in the yeast *Saccharomyces cerevisiae*. Genetics 144, 71–86 (1996).

15. Xu, L., Weiner, B.M. & Kleckner, N. Meiotic cells monitor the status of the interhomolog recombination complex. Genes Dev 11, 106–18 (1997).

16. Sun, X. et al. Transcription dynamically patterns the meiotic chromosome-axis interface. Elife 4, e07424 (2015).

17. Allers, T. & Lichten, M. Differential timing and control of noncrossover and crossover recombination during meiosis. Cell 106, 47–57 (2001).

18. Argunhan, B. et al. Direct and indirect control of the initiation of meiotic recombination by DNA damage checkpoint mechanisms in budding yeast. PLoS One 8, e65875 (2013).

19. Blitzblau, H.G. & Hochwagen, A. ATR/Mec1 prevents lethal meiotic recombination initiation on partially replicated chromosomes in budding yeast. Elife 2, e00844 (2013).

20. Gray, S., Allison, R.M., Garcia, V., Goldman, A.S. & Neale, M.J. Positive regulation of meiotic DNA double-strand break formation by activation of the DNA damage checkpoint kinase Mec1(ATR). Open Biol 3, 130019 (2013).

21. Miyoshi, T. et al. A central coupler for recombination initiation linking chromosome architecture to S phase checkpoint. Mol Cell 47, 722–33 (2012).

22. Murakami, H. & Keeney, S. Temporospatial coordination of meiotic DNA replication and recombination via DDK recruitment to replisomes. Cell 158, 861–873 (2014).

23. Ogino, K. & Masai, H. Rad3-Cds1 mediates coupling of initiation of meiotic recombination with DNA replication. Mei4-dependent transcription as a potential target of meiotic checkpoint. J Biol Chem 281, 1338–44 (2006).

24. Rockmill, B. et al. High throughput sequencing reveals alterations in the recombination signatures with diminishing Spo11 activity. PLoS Genet 9, e1003932 (2013).

25. Thacker, D., Mohibullah, N., Zhu, X. & Keeney, S. Homologue engagement controls meiotic DNA break number and distribution. Nature 510, 241–6 (2014).

26. Tonami, Y., Murakami, H., Shirahige, K. & Nakanishi, M. A checkpoint control linking meiotic S phase and recombination initiation in fission yeast. Proc Natl Acad Sci U S A 102, 5797–801 (2005).

27. Carballo, J.A. et al. Budding yeast ATM/ATR control meiotic double-strand break (DSB) levels by down-regulating Rec114, an essential component of the DSB-machinery. PLoS Genet 9, e1003545 (2013).

28. Garcia, V., Gray, S., Allison, R.M., Cooper, T.J. & Neale, M.J. Tel1(ATM)-mediated interference suppresses clustered meiotic double-strand-break formation. Nature 520, 114–8 (2015).

29. Lange, J. et al. ATM controls meiotic double-strand-break formation. Nature 479, 237–40 (2011).

30. Mohibullah, N. & Keeney, S. Numerical and spatial patterning of yeast meiotic DNA breaks by Tel1. Genome Res 27, 278–288 (2017).

31. Zhang, L., Kim, K.P., Kleckner, N.E. & Storlazzi, A. Meiotic double-strand breaks occur once per pair of (sister) chromatids and, via Mec1/ATR and Tel1/ATM, once per quartet of chromatids. Proc Natl Acad Sci U S A 108, 20036–41 (2011).

32. Kauppi, L. et al. Numerical constraints and feedback control of double-strand breaks in mouse meiosis. Genes Dev 27, 873–86 (2013).

33. Subramanian, V.V. et al. Chromosome Synapsis Alleviates Mek1-Dependent Suppression of Meiotic DNA Repair. PLoS Biol 14 e1002369 (2016).

34. Wojtasz, L. et al. Mouse HORMAD1 and HORMAD2, two conserved meiotic chromosomal proteins, are depleted from synapsed chromosome axes with the help of TRIP13 AAA-ATPase. PLoS Genet 5, e1000702 (2009).

35. Rosu, S. et al. The C. elegans DSB-2 protein reveals a regulatory network that controls competence for meiotic DSB formation and promotes crossover assurance. PLoS Genet 9, e1003674 (2013).

36. Stamper, E.L. et al. Identification of DSB-1, a protein required for initiation of meiotic recombination in Caenorhabditis elegans, illuminates a crossover assurance checkpoint. PLoS Genet 9, e1003679 (2013).

37. Borner, G.V., Barot, A. & Kleckner, N. Yeast Pch2 promotes domainal axis organization, timely recombination progression, and arrest of defective recombinosomes during meiosis. Proc Natl Acad Sci U S A 105, 3327–32 (2008).

38. San-Segundo, P.A. & Roeder, G.S. Pch2 links chromatin silencing to meiotic checkpoint control. Cell 97, 313–24 (1999).

39. Herruzo, E. et al. The Pch2 AAA+ ATPase promotes phosphorylation of the Hop1 meiotic checkpoint adaptor in response to synaptonemal complex defects. Nucleic Acids Res 44, 7722–41 (2016).

40. Roig, I. et al. Mouse TRIP13/PCH2 is required for recombination and normal higher-order chromosome structure during meiosis. PLoS Genet 6, e1001062 (2010).

41. Xu, L., Ajimura, M., Padmore, R., Klein, C. & Kleckner, N. *NDT80*, a meiosis-specific gene required for exit from pachytene in *Saccharomyces cerevisiae*. Mol Cell Biol 15, 6572–81 (1995).

42. Backstrom, N. et al. The recombination landscape of the zebra finch *Taeniopygia guttata* genome. Genome Res 20, 485–95 (2010).

43. Chakraborty, P. et al. Modulating Crossover Frequency and Interference for Obligate Crossovers in *Saccharomyces cerevisiae* Meiosis. G3 (Bethesda) 7, 1511–1524 (2017).

44. Kaback, D.B. Chromosome-size dependent control of meiotic recombination in humans. Nat Genet 13, 20–1 (1996).

45. Kaback, D.B., Guacci, V., Barber, D. & Mahon, J.W. Chromosome size-dependent control of meiotic recombination. Science 256, 228–32 (1992).

46. Kaback, D.B., Steensma, H.Y. & de Jonge, P. Enhanced meiotic recombination on the smallest chromosome of *Saccharomyces cerevisiae*. Proc Natl Acad Sci U S A 86, 3694–8 (1989).

47. Blitzblau, H.G., Bell, G.W., Rodriguez, J., Bell, S.P. & Hochwagen, A. Mapping of meiotic single-stranded DNA reveals double-stranded-break hotspots near centromeres and telomeres. Curr Biol 17, 2003–12 (2007).

48. Gerton, J.L. et al. Global mapping of meiotic recombination hotspots and coldspots in the yeast *Saccharomyces cerevisiae*. Proc Natl Acad Sci U S A 97, 11383–90 (2000).

49. Lam, I. & Keeney, S. Nonparadoxical evolutionary stability of the recombination initiation landscape in yeast. Science 350, 932–7 (2015).

50. Pan, J. et al. A hierarchical combination of factors shapes the genome-wide topography of yeast meiotic recombination initiation. Cell 144, 719–31 (2011).

51. Fung, J.C., Rockmill, B., Odell, M. & Roeder, G.S. Imposition of crossover interference through the nonrandom distribution of synapsis initiation complexes. Cell 116, 795–802 (2004).

52. Serrentino, M.E., Chaplais, E., Sommermeyer, V. & Borde, V. Differential association of the conserved SUMO ligase Zip3 with meiotic double-strand break sites reveals regional variations in the outcome of meiotic recombination. PLoS Genet 9, e1003416 (2013).

53. Chu, S. & Herskowitz, I. Gametogenesis in yeast is regulated by a transcriptional cascade dependent on Ndt80. Mol Cell 1, 685–96 (1998).

54. Prieler, S., Penkner, A., Borde, V. & Klein, F. The control of Spo11’s interaction with meiotic recombination hotspots. Genes Dev 19, 255–69 (2005).

55. Shroff, R. et al. Distribution and dynamics of chromatin modification induced by a defined DNA double-strand break. Curr Biol 14, 1703–11 (2004).

56. Unal, E. et al. DNA damage response pathway uses histone modification to assemble a double-strand break-specific cohesin domain. Mol Cell 16, 991–1002 (2004).

57. Kim, J.A., Kruhlak, M., Dotiwala, F., Nussenzweig, A. & Haber, J.E. Heterochromatin is refractory to gamma-H2AX modification in yeast and mammals. J Cell Biol 178, 209–18 (2007).

58. Szilard, R.K. et al. Systematic identification of fragile sites via genome-wide location analysis of gamma-H2AX. Nat Struct Mol Biol 17, 299–305 (2010).

59. Mimitou, E.P., Yamada, S. & Keeney, S. A global view of meiotic double-strand break end resection. Science 355, 40–45 (2017).

60. Kugou, K. et al. Rec8 guides canonical Spo11 distribution along yeast meiotic chromosomes. Mol Biol Cell 20, 3064–76 (2009).

61. Carballo, J.A., Johnson, A.L., Sedgwick, S.G. & Cha, R.S. Phosphorylation of the axial element protein Hop1 by Mec1/Tel1 ensures meiotic interhomolog recombination. Cell 132, 758–70 (2008).

62. Niu, H. et al. Partner choice during meiosis is regulated by Hop1-promoted dimerization of Mek1. Mol Biol Cell 16, 5804–18 (2005).

63. Vader, G. et al. Protection of repetitive DNA borders from self-induced meiotic instability. Nature 477, 115–9 (2011).

64. Smith, A.V. & Roeder, G.S. The yeast Red1 protein localizes to the cores of meiotic chromosomes. J Cell Biol 136, 957–67 (1997).

65. Conrad, M.N., Dominguez, A.M. & Dresser, M.E. Ndj1p, a meiotic telomere protein required for normal chromosome synapsis and segregation in yeast. Science 276, 1252–5 (1997).

66. Chu, D.B., Gromova, T., Newman, T.A.C. & Burgess, S.M. The Nucleoporin Nup2 Contains a Meiotic-Autonomous Region that Promotes the Dynamic Chromosome Events of Meiosis. Genetics 206, 1319–1337 (2017).

67. Niu, H. et al. Mek1 kinase is regulated to suppress double-strand break repair between sister chromatids during budding yeast meiosis. Mol Cell Biol 27, 5456–67 (2007).

68. Woltering, D. et al. Meiotic segregation, synapsis, and recombination checkpoint functions require physical interaction between the chromosomal proteins Red1p and Hop1p. Mol Cell Biol 20, 6646–58 (2000).

69. Hayashi, M., Mlynarczyk-Evans, S. & Villeneuve, A.M. The synaptonemal complex shapes the crossover landscape through cooperative assembly, crossover promotion and crossover inhibition during *Caenorhabditis elegans* meiosis. Genetics 186, 45–58 (2010).

70. Libuda, D.E., Uzawa, S., Meyer, B.J. & Villeneuve, A.M. Meiotic chromosome structures constrain and respond to designation of crossover sites. Nature 502, 703–706 (2013).

71. Nadarajan, S. et al. Polo-like kinase-dependent phosphorylation of the synaptonemal complex protein SYP-4 regulates double-strand break formation through a negative feedback loop. Elife 6, e23437 (2017).

72. Rog, O., Kohler, S. & Dernburg, A.F. The synaptonemal complex has liquid crystalline properties and spatially regulates meiotic recombination factors. Elife 6, e21455 (2017).

73. Mitra, N. & Roeder, G.S. A novel nonnull *ZIP1* allele triggers meiotic arrest with synapsed chromosomes in *Saccharomyces cerevisiae*. Genetics 176, 773–87 (2007).

74. Mekhail, K. & Moazed, D. The nuclear envelope in genome organization, expression and stability. Nat Rev Mol Cell Biol 11, 317–28 (2010).

75. Srikumar, T., Lewicki, M.C. & Raught, B. A global S. cerevisiae small ubiquitin-related modifier (SUMO) system interactome. Mol Syst Biol 9, 668 (2013).

76. Hayashi, A., Ogawa, H., Kohno, K., Gasser, S.M. & Hiraoka, Y. Meiotic behaviours of chromosomes and microtubules in budding yeast: relocalization of centromeres and telomeres during meiotic prophase. Genes Cells 3, 587–601 (1998).

77. Jin, Q.W., Fuchs, J. & Loidl, J. Centromere clustering is a major determinant of yeast interphase nuclear organization. J Cell Sci 113 (Pt 11), 1903–12 (2000).

78. Wu, H.Y., Ho, H.C. & Burgess, S.M. Mek1 kinase governs outcomes of meiotic recombination and the checkpoint response. Curr Biol 20, 1707–16 (2010).

79. Chen, S.Y. et al. Global analysis of the meiotic crossover landscape. Dev Cell 15, 401–15 (2008).

80. Vincenten, N. et al. The kinetochore prevents centromere-proximal crossover recombination during meiosis. Elife 4(2015).

81. Hoggard, T.A. et al. Yeast heterochromatin regulators Sir2 and Sir3 act directly at euchromatic DNA replication origins. PLoS Genet 14, e1007418 (2018).

82. Schmid, M. et al. Nup-PI: the nucleopore-promoter interaction of genes in yeast. Mol Cell 21, 379–91 (2006).

83. Dilworth, D.J. et al. The mobile nucleoporin Nup2p and chromatin-bound Prp20p function in endogenous NPC-mediated transcriptional control. J Cell Biol 171, 955–65 (2005).

84. Kim, S. et al. The dynamic three-dimensional organization of the diploid yeast genome. Elife 6, e23623 (2017).

85. Ishii, K., Arib, G., Lin, C., Van Houwe, G. & Laemmli, U.K. Chromatin boundaries in budding yeast: the nuclear pore connection. Cell 109, 551–62 (2002).

86. Lange, J. et al. The Landscape of Mouse Meiotic Double-Strand Break Formation, Processing, and Repair. Cell 167, 695–708 (2016).

87. Zanders, S., Sonntag Brown, M., Chen, C. & Alani, E. Pch2 modulates chromatid partner choice during meiotic double-strand break repair in Saccharomyces cerevisiae. Genetics 188, 511–21 (2011).

88. Lee, C.Y., Conrad, M.N. & Dresser, M.E. Meiotic chromosome pairing is promoted by telomere-led chromosome movements independent of bouquet formation. PLoS Genet 8, e1002730 (2012).

89. Auton, A. et al. A fine-scale chimpanzee genetic map from population sequencing. Science 336, 193–8 (2012).

90. Barnes, T.M., Kohara, Y., Coulson, A. & Hekimi, S. Meiotic recombination, noncoding DNA and genomic organization in *Caenorhabditis elegans*. Genetics 141, 159–79 (1995).

91. Mohrenweiser, H.W., Tsujimoto, S., Gordon, L. & Olsen, A.S. Regions of sex-specific hypo- and hyper-recombination identified through integration of 180 genetic markers into the metric physical map of human chromosome 19. Genomics 47, 153–62 (1998).

92. Rockman, M.V. & Kruglyak, L. Recombinational landscape and population genomics of *Caenorhabditis elegans*. PLoS Genet 5, e1000419 (2009).

93. Singhal, S. et al. Stable recombination hotspots in birds. Science 350, 928–32 (2015).

94. Yu, A. et al. Comparison of human genetic and sequence-based physical maps. Nature 409, 951–3 (2001).

95. Kochakpour, N. & Moens, P.B. Sex-specific crossover patterns in Zebrafish (*Danio rerio*). Heredity (Edinb) 100, 489–95 (2008).

96. Zhu, X. & Keeney, S. High-Resolution Global Analysis of the Influences of Bas1 and Ino4 Transcription Factors on Meiotic DNA Break Distributions in *Saccharomyces cerevisiae*. Genetics 201, 525–42 (2015).

97. Blitzblau, H.G., Chan, C.S., Hochwagen, A. & Bell, S.P. Separation of DNA replication from the assembly of break-competent meiotic chromosomes. PLoS Genet 8, e1002643 (2012).

98. Paul, M.R., Markowitz, T.E., Hochwagen, A. & Ercan, S. Condensin Depletion Causes Genome Decompaction Without Altering the Level of Global Gene Expression in *Saccharomyces cerevisiae*. Genetics (2018).

99. Yue, J.X. et al. Contrasting evolutionary genome dynamics between domesticated and wild yeasts. Nat Genet 49, 913–924 (2017).

100. Langmead, B., Trapnell, C., Pop, M. & Salzberg, S.L. Ultrafast and memory-efficient alignment of short DNA sequences to the human genome. Genome Biol 10, R25 (2009).

